# Combinatorial mutagenesis of rapidly-evolving residues yields super-restrictor antiviral proteins

**DOI:** 10.1101/557264

**Authors:** Rossana Colón-Thillet, Emily Hsieh, Laura Graf, Richard N McLaughlin, Janet M. Young, Georg Kochs, Michael Emerman, Harmit S. Malik

## Abstract

Antagonistic interactions drive host-virus evolutionary arms-races, which often manifest as recurrent amino acid changes (*i.e*., positive selection) at their protein-protein interaction interfaces. Here, we investigated whether combinatorial mutagenesis of positions under positive selection in a host antiviral protein could enhance its restrictive properties. We tested ~700 variants of human MxA, generated by combinatorial mutagenesis, for their ability to restrict Thogoto orthomyxovirus (THOV). We identified MxA super-restrictors with increased binding to THOV NP target protein and 10-fold higher anti-THOV restriction relative to wild-type human MxA, the most potent naturally-occurring anti-THOV restrictor identified. Our findings reveal a means to elicit super-restrictor antiviral proteins by leveraging signatures of positive selection. Although some MxA super-restrictors of THOV were impaired in their restriction of H5N1 influenza A virus (IAV), other super-restrictor variants increased THOV restriction without impairment of IAV restriction. Thus, broadly acting antiviral proteins such as MxA mitigate breadth-versus-specificity tradeoffs that could otherwise constrain their adaptive landscape.

## Introduction

The innate arm of mammalian immunity includes dozens of antiviral proteins that act cell-autonomously to block viral replication [1, 2]. Over long evolutionary periods, innate immune proteins must adapt to combat rapidly-evolving pathogenic viruses. This adaptation often occurs at the interfaces between viral and host immune proteins, to either evade or increase binding interactions [3]. This back-and-forth evolution between host and viral proteins can result in signatures of positive selection (higher rates of non-synonymous substitutions, dN, compared to synonymous substitutions, dS) that can be identified by a comparison of orthologous genes over a phylogenetic tree. Positive selection is often concentrated in just a few residues that maximally impact binding affinity and thereby dictate the outcomes of viral infection in host cells [3]. Indeed, experimental tests show that variation in positively selected sites can explain the species-specific differences in antiviral specificity between orthologous antiviral proteins [3–5].

Such retrospective analyses are a powerful means of identifying amino acids and protein-protein interaction interfaces that have been shaped by past episodes of selection. Nonetheless, they are limited in terms of describing the landscape of possible adaptation against viruses. For example, epistasis between different rapidly evolving sites could have shaped and constrained the available landscape of possible adaptation against viruses [6–8]. We sought to leverage the previous insights gained from our retrospective analyses to design a prospective means to elicit higher-specificity versions of antiviral proteins and understand the available paths to the adaptation of a host innate immune protein to potential viral pathogens.

We focused our studies on MxA, an interferon-induced large dynamin-like GTPase, which inhibits a diverse range of viruses by interacting with multiple distinct viral proteins [9, 10]. MxA comprises an N-terminal globular GTPase-containing head (G domain) and a C-terminal stalk, which are connected by a hinge-like bundle-signaling element (BSE) [11, 12]. Previously, we had identified the L4 loop (L4), which protrudes from the MxA C-terminal stalk, as a hotspot for recurrent positive selection in primates [4]. We showed that five residues (amino acids 540, 561, 564, 566, 567 in human MxA) in L4 have evolved under positive selection during simian primate evolution. Moreover, single amino acid changes at amino acid 561 in the L4 region of MxA were necessary and sufficient to explain dramatic differences in the species-specific antiviral activity of primate MxA proteins against Thogoto virus (THOV) and influenza A virus (IAV), two distantly related orthomyxoviruses [4].

We hypothesized that combinatorial mutagenesis of all five positively selected residues in L4 of human MxA might reveal the contributions of positions other than residue 561 in antiviral restriction and generate MxA variants with increased antiviral activity (here, called ‘super-restrictors’) relative to human MxA. Our combinatorial analyses revealed strict amino acid requirements at some L4 positions (*e.g*., residue 561) but also significant contributions of other L4 positions (*e.g*., residues 540, 564) to the gain and loss of restriction against THOV. Our analyses also revealed epistasis between L4 residues, thereby reiterating the merit of our combinatorial mutagenesis approach. Finally, consistent with our predictions, our studies recover super-restrictor variants of human MxA that have 10-fold higher antiviral activity against THOV, which correlates with increased binding to the THOV nucleoprotein (NP). Finally, we find that MxA super-restriction against one virus is not accompanied by its increased restriction against a second, related viral target. Although our strongest MxA super-restrictors against THOV lost activity against another orthomyxovirus, H5N1 IAV, a few THOV super-restrictors retained full IAV restriction activity. Our analyses thus reveal not only a powerful means to elicit super-restrictor versions of antiviral proteins, but also provide insights into how antiviral proteins like MxA may skirt breadth-versus-specificity tradeoffs that might otherwise constrain their adaptation against multiple viruses.

## Results

### Combinatorial mutagenesis reveals L4 sequence requirements for THOV restriction

Retrospective evolutionary analyses are a powerful means to decipher the landscape of selective constraint on protein-coding genes. Such studies can not only identify which residues have recurrently evolved due to relentless adaptation (positive selection) [3], but also identify residues that have not undergone any changes despite millions of years of protein divergence (purifying selection). We reasoned that a prospective approach focusing on combinatorial mutagenesis of positively selected sites, while preserving sites that have evolved under purifying selection, might reveal insights into the selective pressures that mold antiviral defense repertoires and uncover unexplored potential for enhanced antiviral activity. Furthermore, we reasoned that focusing on only a subset of sites revealed to be critical for past adaptation by retrospective studies would allow us to use combinatorial mutagenesis to survey potential paths of antiviral adaptation that involved changes at more than one site.

We, therefore, generated a *MxA* gene variant library (Fig 1A, S1_Fig) encoding MxA proteins containing random combinations of all amino acids in the five positively selected sites of L4 in wild-type human MxA (hereafter referred to as wtMxA). Five residues in the L4 of wtMxA protein have evolved under recurrent positive selection in simian primates: 540 (G), 561 (F), 564 (F), 566 (S) and 567 (S). These residues were combinatorially mutagenized to all 20 amino acids using oligos with degenerate NNS codons (Materials and Methods). We randomly selected 523 clones (discarding those that had stop codons or mutations outside L4) and individually evaluated their antiviral activity against THOV using a minireplicon assay (Materials and Methods, S2_Fig). In this minireplicon assay, the viral polymerase components (PB2, PB1, PA), as well as the genomic-RNA-binding nucleoprotein (NP) drive the expression of a firefly luciferase reporter. Overexpression of wtMxA reduces THOV-mediated luciferase expression in a dose-responsive manner, while a mutation in the MxA GTPase catalytic domain, T103A, renders human MxA non-restrictive against THOV regardless of MxA plasmid input concentration (S2_Fig) [4]. By measuring the restrictive activities of T103A and wtMxA multiple times, we determined the range of variation of wtMxA THOV restriction activity (Fig S3) and then compared the restriction of all MxA variants (Fig 1B, S1_Table).

**Figure 1.**
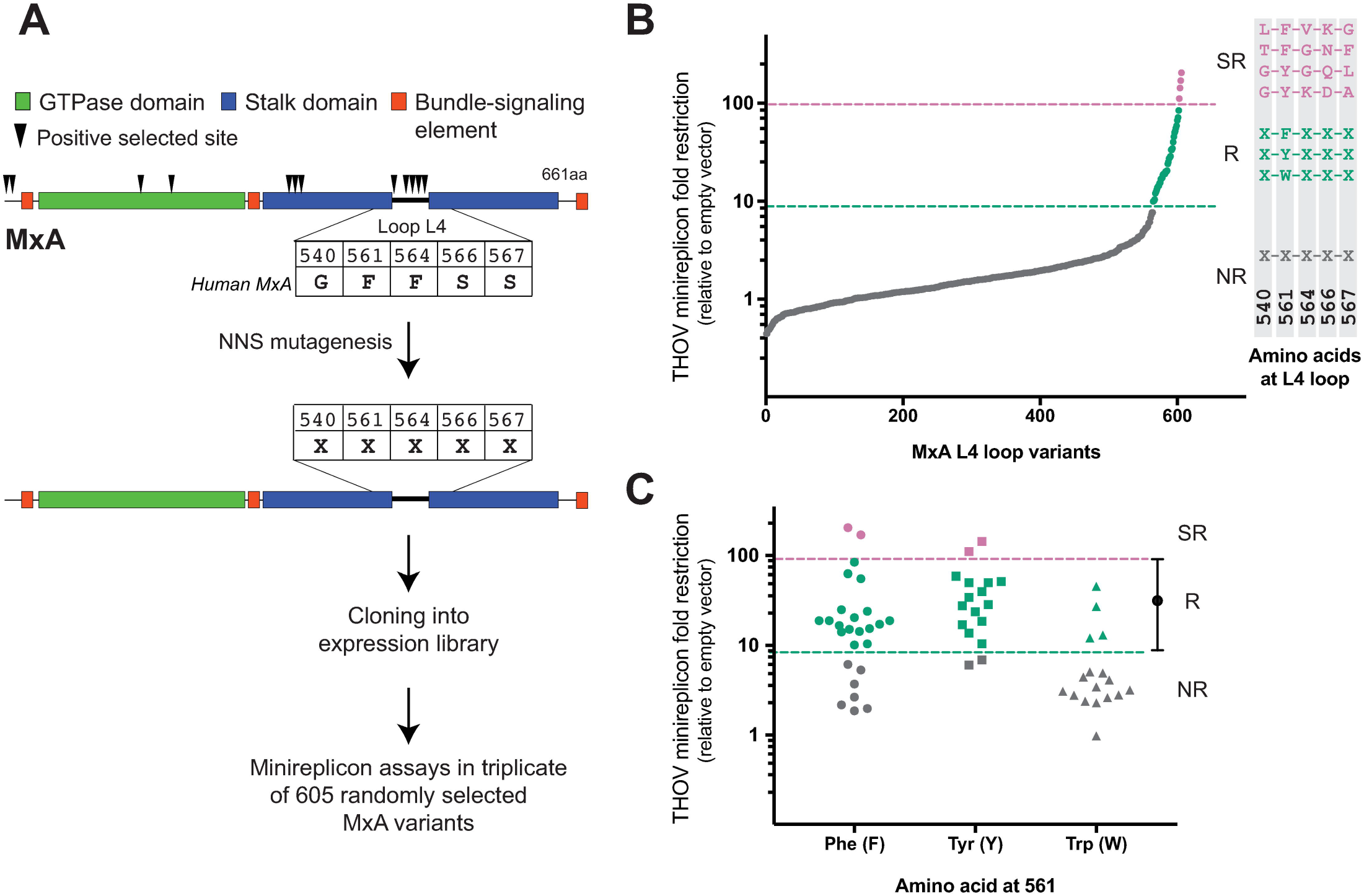
Combinatorial random mutagenesis of rapidly evolving sites in the MxA L4 loop alters antiviral potency against THOV. **A.** A positive selection-based mutagenesis scheme to obtain MxA combinatorial variants of the 5 rapidly-evolving sites (arrowheads) in L4 (N = any nucleotide, S = G or C). **B.** Random MxA variants are presented in order of increasing THOV fold restriction, with their sequence features summarized to the right. ‘Non-restrictor’(NR) variants with lower restriction than wtMxA are shown below the dashed green line (these contain very diverse residues in the randomized L4 positions). Approximately 6% of the variants have equivalent restriction to wtMxA (green restrictors or ‘R’, between the pink and green dashed lines) and these all possess a F, Y, or W residue at position 561. L4 sequences of four variants that restrict THOV better than wtMxA (pink super-restrictors or ‘SR’) are shown. **C.** Fold restriction of MxA variants recovered with F, Y, or W residues at position 561. All data points represent averages of minireplicon experiments done in triplicate for each variant.

Despite only altering the five most rapidly evolving L4 positions, we found that most MxA variants (~95%) had worse THOV restriction than wtMxA, consistent with the evolutionary finding that these residues are crucial for MxA’s antiviral function. We searched for patterns to explain the sequence requirements to maintain MxA antiviral activity against THOV. We found that all restrictive variants possessed hydrophobic aromatic amino acids (Phe F, Tyr Y or Trp W) at residue 561 (Fig 1B, right; S1_Table). These results extend our previous findings that a hydrophobic aromatic amino acid at residue 561 could confer antiviral restriction against THOV in the context of wtMxA [4]. While many non-restrictive variants had non-aromatic residues at position 561, we obtained at least 21 combinatorial MxA variants that had weaker anti-THOV activity than wtMxA despite possessing F, Y, or W at residue 561 (Fig 1C, grey shapes). Therefore, we conclude that an aromatic residue at position 561 is necessary but not sufficient to confer anti-THOV restriction. Our findings imply that the other four rapidly evolving L4 residues also directly contribute to MxA antiviral activity against THOV.

### Combinatorial mutagenesis of positively selected residues yields super-restrictor variants

In our initial screen, we also discovered four MxA variants (LFVKG, GYKDA, TFGNF and GYGQL in the randomized L4 positions) with 2-to 5-fold better restrictive ability against THOV than wtMxA (Figs 1B and 1C in pink) in the replicon assay. This increased potency cannot be explained by expression levels (Fig 2B, western blot panel). These super-restrictor MxA variants validate our premise that combinatorial mutagenesis of positively selected residues could yield increase antiviral potency. However, only 6% of the initial mutant pool possessed the aromatic amino acid at position 561 (Fig 1B in green and pink, Fig 1C) required for THOV restriction. Therefore, we designed a second combinatorial library, in which we fixed site 561 as F, the amino acid present in wtMxA while randomizing the other four amino acids under positive selection in L4.

**Figure 2.**
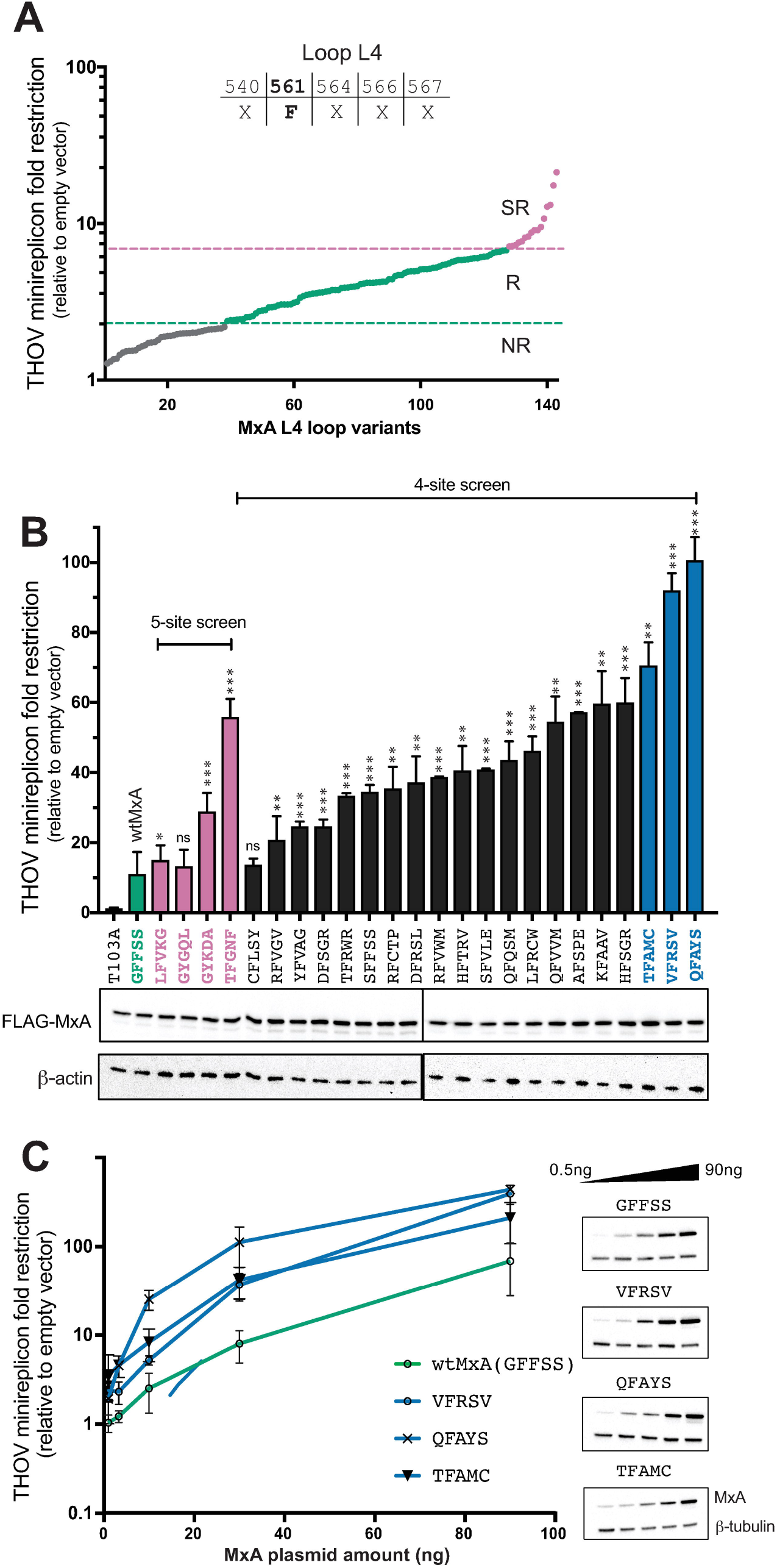
A modified mutagenesis strategy yields potent THOV super-restrictor variants. **A.** A modified combinatorial mutagenesis strategy with fixed F561 and four randomized L4 positions. Non-restrictor, restrictor, and super-restrictor variants are presented in order of increasing restriction activity (X-axis). **B.** Re-testing of anti-THOV restriction activity of super-restrictor MxA variants (relative to the empty vector) obtained from the original five-site screen (pink) and the modified 4-site screen (black). The three most potent super-restrictors (blue), have 7-to 10-fold higher anti-THOV restriction activity relative to wtMxA. Data is presented as mean of three experiments (error bars ±SEM). For super-restrictor variants, restriction values were compared to the wtMxA control in the experiment *p ≤ 0.05, **p ≤ 0.01, ***p ≤ 0.001, ****p ≤ 0.0001 (Mann-Whitney U test). Western blots indicate levels of expression of transfected MxA variants relative to a loading control (β-actin). **C.** Dose-responsiveness of super-restriction activity for MxA variants. Western blots indicate levels of expression of each of transfected MxA variants at various plasmid concentrations (top row) relative to a loading control (□-tubulin, bottom row). The data underlying this figure can be found in S1_Data.

For the four-site screen, we randomly selected 141 variants and assessed their antiviral activity in triplicate against THOV using the minireplicon assay (Fig 2A, S2_Table). As predicted, a higher proportion (~65%) of MxA variants in the 4-site screen have anti-THOV activity comparable to wtMxA (Fig 2A, S2_Table) as opposed to only ~6% in the five-site screen (Fig 1B). Moreover, we found that ~13% of the 4-site variants are super-restrictors (Fig 2A) compared to only ~0.5% in the initial screen (Fig 1B). To validate the top super-restrictors from both screens, we re-tested their activity in the THOV minireplicon assay (Fig 2B). Most of the variants identified as super-restrictors again showed enhanced restriction of THOV. In particular, the three most potent super-restrictors obtained with the 4-site screen (*i.e*., QFAYS, VFRSV, and TFAMC) are 7 to 10-fold more potent than wtMxA at restricting THOV in the minireplicon assay (Fig 2B). Although these super-restrictor variants are expressed at similar levels as wtMxA (Fig 2C, bottom), they have higher anti-THOV restriction at all levels of expression (Fig 2C). This significant improvement in THOV restriction is remarkable because human wtMxA is the most potent naturally-occurring MxA ortholog for THOV restriction identified to date [4, 13].

To confirm the results of the minireplicon assay, we also compared the restriction of wtMxA, two non-restrictor variants (PFFSS and ΔL4), and the three strongest MxA super-restrictors, in infection experiments where Huh7 cells were challenged with replication-competent THOV virus (Fig 3A). Despite the lower dynamic range of this assay compared to the minireplicon experiments, we found that two super-restrictor MxA variants (TFAMC and QFAYS) restricted THOV infection significantly better than wtMxA whereas increased restriction was not statistically significant for a third variant (VFRSV).

**Figure 3.**
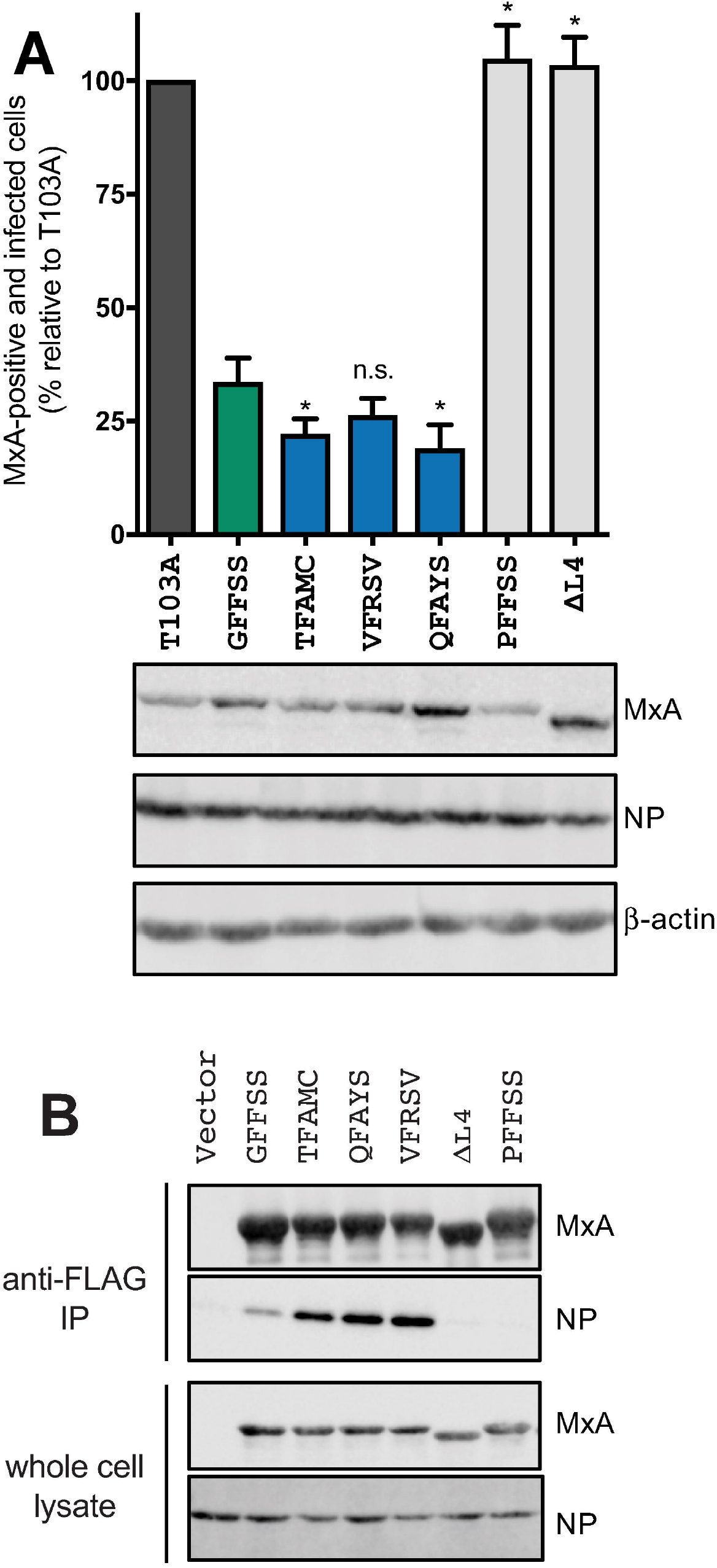
Restriction and biochemical properties of MxA variants in THOV-infected cells. **A.** Using THOV infection of Huh7 cells expressing different MxA variants, we compared THOV restriction provided by wtMxA, MxA super-restrictors or MxA non-restrictors (PFFSS or ΔL4) relative to a catalytically inactive MxA (T103A) mutant. Statistical significance of increased restriction of super-restrictors relative to wtMxA was ascertained using a 1-way ANOVA test (*p ≤ 0.05). Western blots show the levels of MxA variants, THOV NP protein and a loading control (β-actin). **B.** Co-immunoprecipitation studies of viral NP target protein with different MxA variants using cell lysates of THOV-infected 293T cells transfected with the FLAG-tagged MxA expression constructs. MxA super-restrictors pull down more THOV NP protein than wtMxA, whereas MxA non-restrictors (PFFSS or ΔL4) pull down significantly lower amounts of THOV NP protein. Western blots of the FLAG-tagged MxA variants and THOV NP protein are shown. The data underlying this figure can be found in S1_Data.

Since we previously found that changes at amino acid 561 correlated with gain or loss of binding to the target nucleoprotein (NP) [4, 14], we next tested if increased MxA-NP interaction could explain the super-restriction phenotype [15–17]. MxA forms higher-order oligomers that are required for antiviral activity [11, 18, 19]. Thus, the increased affinity of MxA monomers to NP might result in higher avidity in the multimeric MxA complex that interacts with THOV NP [4, 11, 14, 18, 20]. We assayed the association of NP with FLAG-tagged MxA super-restrictors in cells infected with THOV. We found that the most potent super-restrictors QFAYS, VFRSV, and TFAMC pull down a larger fraction of cellular NP compared to wtMxA (GFFSS) (Fig 3B), whereas non-restrictor MxA versions (PFFSS and ΔL4) have significantly reduced ability to pull down THOV NP. These data suggest that the mechanism for super-restriction involves a more avid interaction of MxA with the THOV NP target protein during viral infection.

### Multiple L4 residues contribute to MxA super-restriction

We next investigated whether there was a common sequence pattern that could explain the enhanced anti-THOV restriction activity of the super-restrictor variants. We noticed that MxA variants with glutamine (Q) at position 540 were enriched among the 4-site super-restrictors (Fig 2B, S4_Fig), including the most potent super-restrictor, QFAYS. Swapping the glutamine (Q) residue at position 540 in QFAYS, QFQSM, and QFVVM to the glycine (G) present in wtMxA led to a significant drop in restriction activity (Fig 4A). This finding suggested that position 540 could play a critical role in determining super restriction. To better explore the contribution of residue 540 to super-restriction, we mutated the glycine at residue 540 (G540) in the wtMxA backbone to every other amino acid (Fig 4B). We found that changing position 540 to Q in wtMxA leads to only a 2-fold improvement in antiviral activity (Figs 4A and 4B), significantly less than the super-restriction of the QFAYS, QFQSM and QFVVM variants. Moreover, although individual substitutions (*e.g*., G540A or G540Q) could increase THOV restriction 2-3 fold, none of the single substitutions at residue 540 achieved the level of super-restriction in the most potent variants (compare Figs 4B and 2B). These results imply that Q540 is necessary but not sufficient to explain super-restriction. Consistent with these findings, although variants QFQSM and QFVSM are both super-restrictors, variant QFLSM is a non-restrictor despite differing only at position 564 (Fig 4C). Our results imply that multiple positively selected L4 residues in addition to residue 540, can enhance or epistatically interfere (*e.g*., QFLSM) with the degree of MxA restriction.

**Figure 4.**
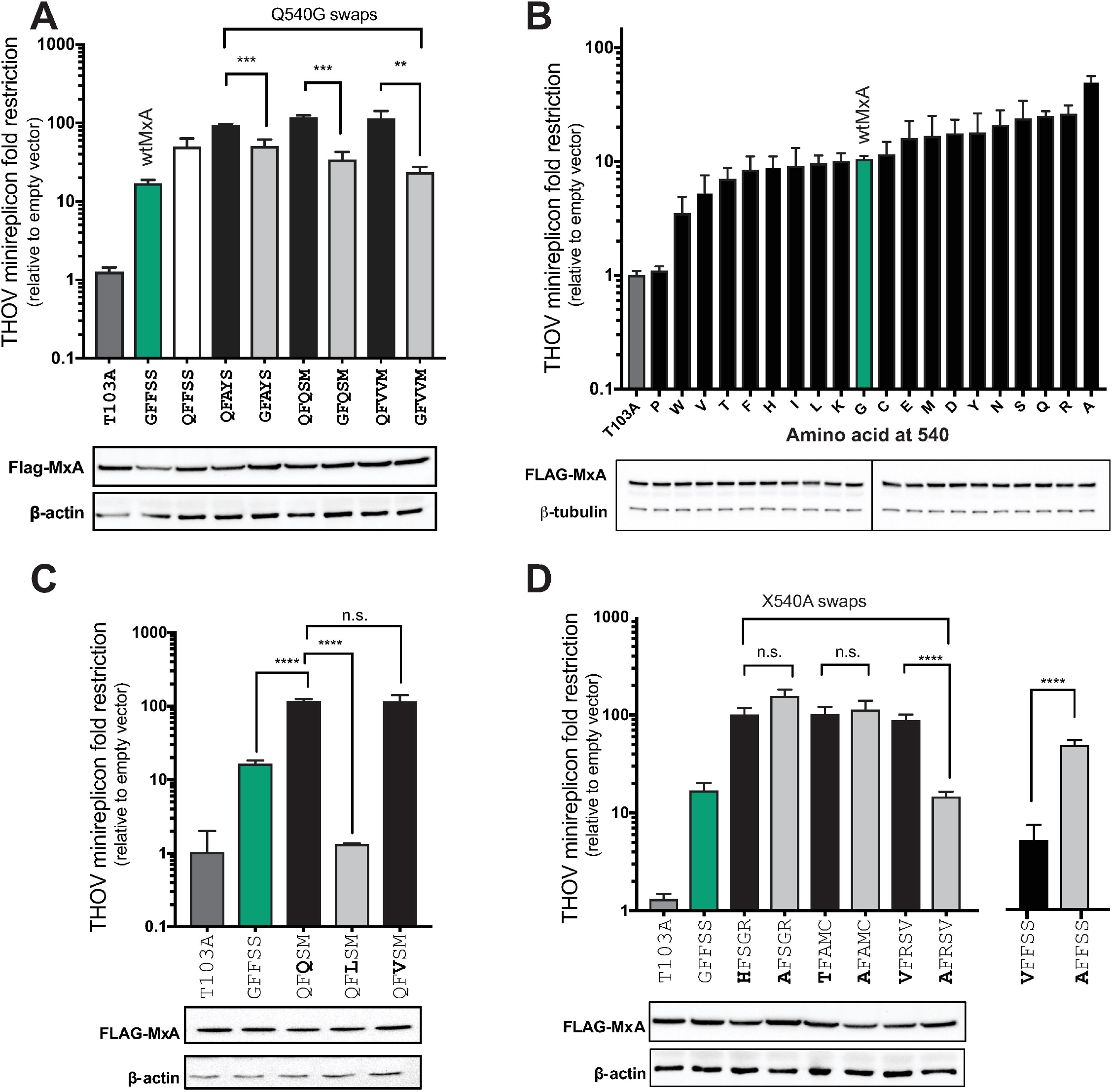
Molecular basis of MxA super-restriction. **A.** Comparison of anti-THOV activity of Q540 (super-restrictor) to Q540G MxA variants. A G540Q mutation in wtMxA resulted in increased restriction (white versus green bar), whereas Q-to-G changes reduced super-restrictor activity of three super-restrictors (black versus gray bars). **B.** Effect of single residue G540X mutations in wtMxA (green) on anti-THOV restriction relative to the empty vector. **C.** Anti-THOV activity of super-restrictor QFQSM MxA variant compared to QFLSM and QFVSM MxA variants demonstrates the contribution of residue 564. **D.** Effect of single residue X540A changes in three MxA super-restrictor variants: HFSGR, TFAMC and VFRSV. All variants are compared to wtMxA *p ≤ 0.05, **p ≤ 0.01, ***p ≤ 0.001, ****p ≤ 0.0001 (Mann-Whitney U test). The data underlying this figure can be found in S1_Data.

Our single residue swap studies revealed 540A to be the most potent restrictor (Fig 4B) in the context of the wtMxA (GFFSS) backbone. However, we speculated that the enhancement of MxA restriction by 540A mutations might also be context-dependent. To test this possibility, we made 540A mutations in three super restrictors – HFSGR, TFAMC, and VFRSV – that each encode for an ‘unfavorable’ amino acid at position 540 in the wtMxA context (Fig 4D). If the increased MxA restriction by 540A were context-independent, we would expect to see further enhancement of anti-THOV activity in each of the HFSGR, TFAMC and VFRSV super-restrictors via 540A mutation. Instead, we found that HFSGR and AFSGR MxA variants were equivalent in their restriction, as were the TFAMC and AFAMC variants (Fig 4D). However, a V540A swap in VFSRV dramatically lowered restriction activity by 10-fold (Fig 4D), even though the same V540A swap in a wtMxA context (XFFSS) increased restriction by 5-fold (Fig 4B). Together, our results show that a simple model of a context-independent contribution of L4 residues cannot account for the broad repertoire of super-restrictors. If our analyses had relied on combining ‘restriction-favorable’ single residues like A540 while avoiding ‘restriction-unfavorable’ residues like V540, we would have never been able to discover super-restrictors like VFRSV. Instead, we find that epistasis shapes the restriction profile of many MxA super-restrictor variants such as VFRSV. This discovery of context-specificity demonstrates the power of an unbiased combinatorial mutation approach rather than a single-residue deep mutational scanning approach to provide a more thorough exploration of the adaptive landscape for discovering novel super-restrictors.

### Breadth-versus-specificity tradeoffs affecting MxA antiviral specificity

Since MxA restricts a number of different viruses, its evolution is likely to have been shaped by a number of host-virus interactions. Therefore, we investigated how restriction against one virus affects the antiviral activity against another virus. Previous studies have shown that restriction of both THOV and H5N1 IAV depends on the L4, specifically on residue F561 in MxA [4]. Since both viruses are restricted by a similar interface, we speculated that gaining super-restriction against THOV via changes in the MxA L4 could also affect (either gain or loss) its restriction of IAV. Alternatively, the gain of THOV restriction could be uncorrelated with IAV restriction.

We, therefore, tested MxA L4 variants against H5N1, an emerging IAV strain that is pathogenic in poultry. We chose the H5N1 strain instead of human IAV strains (*e.g*., H1N1 or H3N2) because human-adapted IAV strains have already evolved evasion of human MxA [16, 17, 21]. We first tested several ‘non-restrictor MxA variants that still preserved an aromatic amino acid at residue 561 but had lost anti-THOV activity (Fig 5A). We tested these MxA variants for their anti-H5N1 IAV restriction activity using the IAV minireplicon assay described previously (Materials and Methods) as a proxy for H5N1 IAV replication. We found that these variants still retained anti-IAV activity (Fig 5B). Thus, loss of THOV restriction by mutations in L4 does not necessitate a loss of H5N1 IAV restriction.

**Figure 5.**
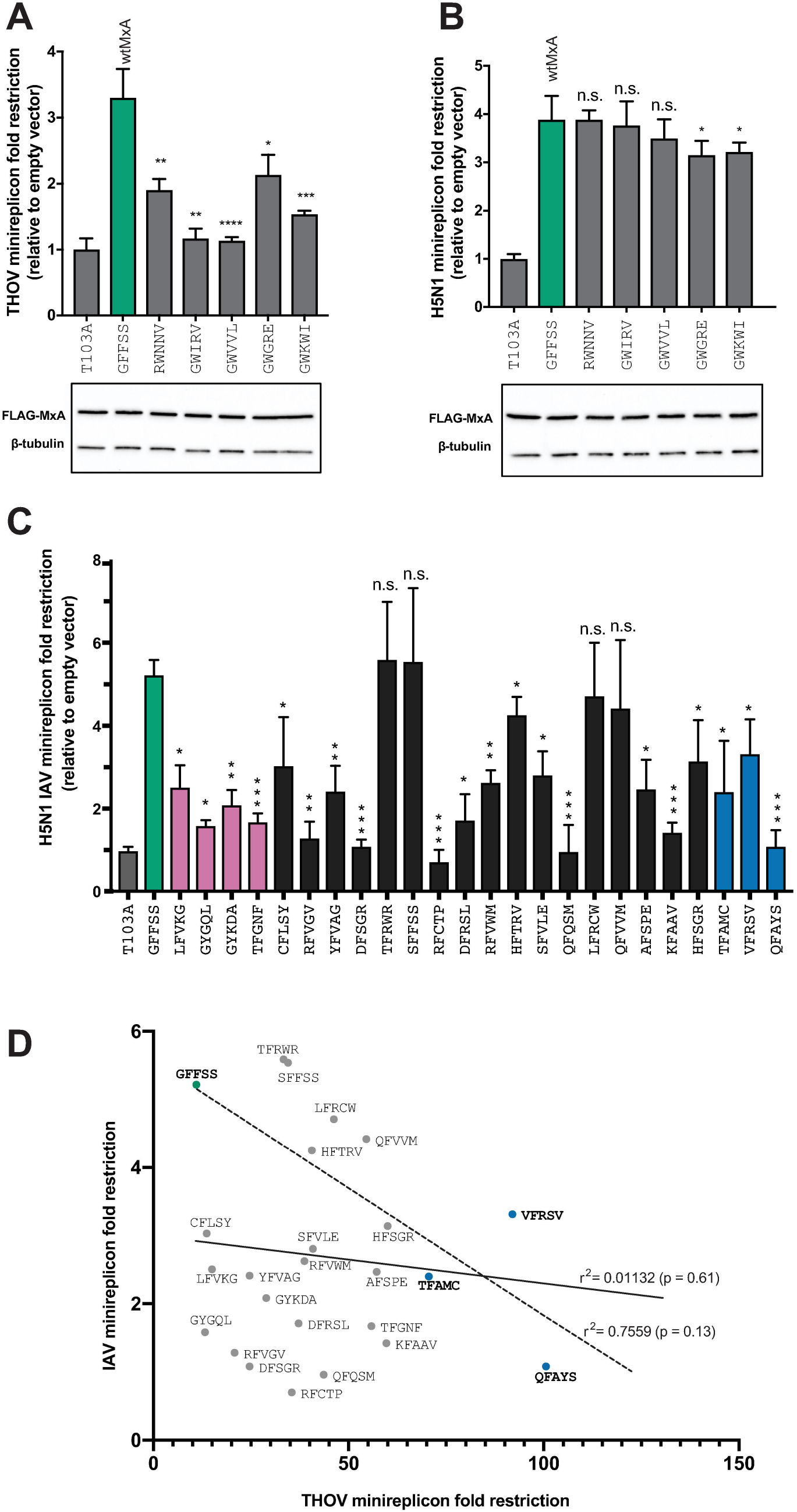
MxA variant restriction of IAV and THOV. **A, B.** MxA non-restrictor’ variants still encoding a necessary aromatic residue at position 561 were re-tested in minireplicon assays to measure restriction against THOV (**A**) and H5N1 IAV (**B**) viruses. Western blots showing the expression levels of FLAG-tagged MxA variants and a loading control (b-tubulin are presented). **C.** Restriction activity of MxA 5-site super-restrictor variants (pink) and 4-site super-restrictor variants (black, blue) against the H5N1 strain of IAV in a minireplicon assay. MxA variants are grouped according to whether they were observed in the 5-site (pink) or 4-site (black, blue) screens and presented in order of increased anti-THOV restriction (from Fig 2B). In all panels, data is presented as means of three experiments (error bars ±SEM) and the IAV restriction of MxA variants is compared to wtMxA based on Mann-Whitney U tests (*p≤0.05, **p ≤ 0.01, ***p ≤ 0.001, **** p ≤ 0.0001). Fig 2B reports the expression levels for various MxA variants. **D.** There is a weak inverse correlation between the THOV and IAV restriction activities for the three most potent anti-THOV super-restrictors VFRSV, TFAMC, and QFAYS compared to wtMxA (Pearson linear correlation r^2^ =0.7559, p=0.13) but this inverse correlation is not observed when comparing all MxA super-restrictors (r2=0.01132, p=0.61). The data underlying this figure can be found in S1_Data.

Next, we tested all 24 MxA super-restrictor variants that we had identified against THOV in both the 5-site (Fig 1) and 4-site screens (Fig 2) against H5N1 in a minireplicon assay (Fig 5C). We found that increase in super-restriction to THOV did not result in a corresponding increase in restriction to IAV (Fig 5C). Thus, gain in super-restriction is virus-specific. Indeed, the three most potent super-restrictors against THOV had significantly lower restriction against THOV (Fig 5C). Similarly, 17 of the other 21 anti-THOV super-restrictors we tested have lower antiviral activity against H5N1 IAV than wtMxA (Fig 5C). On the other hand, four MxA super-restrictor variants (TFRWR, SFFSS, LFRCW, QFVVM) have statistically equivalent restriction of H5N1 IAV as wtMxA (Fig 5C). Comparison of the anti-THOV and anti-IAV restriction activities of the three most potent anti-THOV MxA super-restrictor variants suggested a negative correlation between THOV and IAV restriction (dotted line, Fig 5D). However, comparison of the full set of 24 super-restrictors revealed no correlation between fold-restriction of THOV and fold-restriction of IAV (solid line, Fig 5D).

Thus, despite varying at only 5 rapidly evolving L4 positions, the MxA variants we have analyzed run an entire gamut of phenotypes, ranging from being effective against IAV but not THOV (Figs 5A and 5B), improved restriction of THOV but worse restriction of IAV (Figs 3B and 5C), or enhanced restriction of THOV without any significant impairment against IAV (Fig 5C). Notably, we found no MxA variants that had significantly increased activity against both THOV and H5N1. Our results demonstrate that optimization of MxA to restrict one virus has either detrimental or neutral effects on its restriction activity against another virus. Our findings not only reveal that wtMxA provides a generalist solution to restrict both THOV and IAV and perhaps a large number of other viruses [10], but also demonstrate that its adaptation against a specific virus or viral protein does not require a loss of prior antiviral restrictive abilities. Thus, MxA is able to bypass the breadth-versus-specificity tradeoffs that would otherwise preclude it from acting as a broadly antiviral protein.

## Discussion

Taking advantage of an evolution-guided approach that combines combinatorial mutagenesis with positive selection analyses, we have been able to identify MxA variants that are 10-fold better anti-THOV restrictors than wtMxA, itself the most potent naturally-occurring MxA ortholog known against THOV [4, 13]. Our analyses reveal that these super-restrictors require a contribution from several positions in L4 including the requirement of an aromatic residue at residue 561. Conversely, we find that ‘unfavorable’ residues can also render MxA variants to become ‘non-restrictors’ against THOV in spite of having an aromatic residue at position 561. Thus, multiple L4 residues that have evolved under recurrent positive selection over primate evolution shape the antiviral specificity of MxA.

Our analyses further reveal that epistasis helps determine the contributions of individual L4 residues in shaping MxA specificity. We note instances where the same amino acid mutation (A540V) can have opposite effects on restriction activity in MxA proteins that only differ at three other L4 residues. Our results suggest that the path to adaptation in MxA is indeed shaped by historical contingency (*i.e*., what MxA sequence was present when adaptation occurred) as well as by epistasis. These findings that are reminiscent of previous work delineating the evolutionary path to adaptation is often shaped by contingency and epistasis [6–8, 22]. Although the focus of this study is on positively selected residues in L4 and orthomyxoviruses, other positively selected residues outside L4 may similarly influence super-restriction and epistasis between different domains to determine MxA specificity against other viral families [4, 10].

Our strategy for the discovery of super-restriction factors preserves protein domains subject to purifying selection and focuses on generating and testing mutations at residues already highlighted by evolutionary analyses as recurrent targets of adaptation [3]. This focus on only a smaller subset of residues allows us to explore outcomes of combinatorial mutagenesis [23] by relaxing the constraints imposed by epistasis and historical contingency. Using this method, we are thus able to obtain super-restrictor MxA versions that would not necessarily have been obtained by combining ‘favorable’ residues identified by a single-residue deep-mutational scan of all L4 positions. Our results are based on a shallow sampling of the sequence space of all possible combinations of amino acid substitutions in the sites of positive selection since the minireplicon assay is not amenable to comprehensive sampling of all possible combinatorial mutations. For example, we sampled 523 out of a possible 3.2 million variants in the original 5-site screen and 141 out of 160,000 possible combinations in the second 4-site screen. Nonetheless, despite shallow sampling, we were able to readily identify super-restrictor variants against THOV and show that their activity is influenced by contributions and epistatic interactions of multiple L4 residues.

The positive selection of MxA L4 may not have been contemporaneously shaped via interactions with NP proteins of orthomyxoviruses like THOV or IAV over simian primate evolution. However, these viruses serve as proxies to elucidate the phenotypic consequences of historic selective events. For example, our analysis of MxA variants derived via combinatorial mutagenesis reveals a breadth-versus-specificity tradeoff in which most, but not all, MxA super-restrictor variants against THOV restriction appear to lose H5N1 IAV restriction. While a more in-depth sampling of the library might uncover MxA variants with higher restriction activity against both viruses, our findings suggest that, under threat by multiple viruses, antiviral genes such as MxA appear to be under evolutionary pressure to harbor more broadly active alleles even at the expense of more potent, specific antiviral activity. This result is reminiscent of a recent study, which reported that generalist (promiscuous) MHC class II alleles are selected for in human populations in response to high pathogen diversity [24]. Notably, although MxA specialization to increase restriction against one virus (*e.g*., THOV) is more likely to impair its effectiveness against another (*e.g*., H5N1 IAV), this is not always the case.

Breadth-specificity tradeoffs have been invoked in the case of cytochrome p450 detoxification genes that protect herbivorous insects from plant counter-defenses [25] and in plant disease resistance (R) genes encoded by the Leucine-Rich-Repeat (LRR) gene family [26]. Furthermore, the breadth-specificity tradeoff we observe bears similarity to the outcome of an artificial selection experiment that made the GroEL chaperone highly specific for one substrate but at the expense of its substrate-binding breadth [27]. Remarkably, critical components of the innate immune defense apparatus may evolve under the same constraints as an intracellular chaperone, sacrificing high avidity of binding to a given viral substrate to maintain a breadth of target binding and thereby antiviral response. Gene duplication and divergence is one commonly adopted means to relieve the constraints of such tradeoffs [3, 28–32]. However, despite recurrent duplication and recombination, mammalian genomes rarely encode more than one Mx paralog that localizes to the same subcellular compartment [33]; the biological basis for this constraint against Mx gene duplication is unknown.

Although MxA is unusual in its antiviral breadth, many antiviral proteins evolve under similar selective regimes to MxA, in which they undergo recurrent positive selection in response to viral pathogens [3]. We propose that by leveraging the signature of positive selection, our evolution-guided approach can similarly increase the potency of other antiviral proteins. In many of these cases (*e.g*., TRIM5, MxB), positive selection appears enriched in unstructured loops [4, 5, 33–36] like MxA L4 [18]. We speculate that unstructured loops like L4 represent an ensemble of many conformations [18] that provide MxA with structural and evolutionary flexibility to adapt to bind distinct viral targets. Under this hypothesis, the majority of MxA super-restrictor variants may be less structurally flexible in L4, trading off increased avidity and restriction of specific viral targets with decreased antiviral range. However, a few of the MxA super-restrictors can skirt this tradeoff. Understanding the structural basis of this tradeoff and why it applies to the majority of, but not all, super-restrictors might reveal the biological forces that shape and constrain the adaptive landscape of antiviral proteins. These studies may also help delineate the biological basis of MxA’s unusually broad antiviral activity [10].

## Materials and Methods

### NNS library construction

The L4 loop five-point variant library was constructed using oligonucleotide-directed mutagenesis of the rapidly evolving sites in MxA. To mutate the five rapidly evolving sites in the L4 loop (positions 540, 561, 564, 566 and 567), two mutagenic oligonucleotides (one sense, one antisense) were synthesized (IDT) that contain sequence complementarity to 70 bp in the region encoding for the rapidly evolving residues. For the targeted positions, the oligonucleotides contain NNS codons (N = A, T, C or G, and S = G or C). This biased randomization results in 32 codons with all 20 amino acids sampled –a significant decrease in library complexity and incidence of stop codons without the loss of amino acid complexity. The maximum complexity of this library is thus 20^5^, or 3.2 million variants. One round of PCR was carried out with either the sense or antisense oligonucleotide and a flanking antisense or sense oligonucleotide. A second round of PCR using a combination of first round products and both flanking primers produced the full-length double stranded product. The full-length PCR product was purified, digested and ligated into the pQXCIP retroviral expression vector. The ligation was purified and eluted in 10 μl dH_2_O, which was used to transform ElectroMAX™ DH10B™ cells (ThermoFisher). 836 colonies were randomly selected and cultured overnight in LB liquid medium containing 100 μg/ml of ampicillin. All variants containing stop codons introduced by NNS mutagenesis were removed and not analyzed further.

The L4 loop four-site variant library was similarly constructed, except that only four L4 positions were randomized, whereas position 561 was fixed as F. This library has a maximum complexity of 20^4^, or 160,000 variants. We evaluated the 141 clones that were subsequently analyzed for 1-site, 2-site, 3-site or 4-site mutations relative to wtMxA (S3_Table). We generated an expected distribution based on the NNS mutagenesis approach. We found that the actual distribution of variants were close to the expected distribution except for a slightly higher incidence of 1-site mutations (5 observed versus 0 expected).

### Sequencing

The library variants and all other MxA clones in this study were sequenced in full-length using standard Sanger sequencing with a set of four primers: F1: 5’-CCG CTG GTG CTG AAA CTG AAG AAA C-3’, R1: 5’-ACC ACC AGA TCA GGC TTC GTC AAG A TTC-3’, F2: 5’-ACA GTA TCG TGG TAG AGA GCT GCC-3’ and R2: 5’-AGC ATG GCC CAG GAG GTG GAC CCC G-3’.

### Plasmids

MxA L4 loop variants were cloned into the NotI/EcoRI sites of the retroviral expression vector pQXCIP-3x flag. Point mutants were generated using the Q5^®^ site-directed mutagenesis kit (New England Biolabs).

### Minireplicon assays

The THOV minireplicon assay was performed in 293T cells in a 96-well format. 4.0 ng each of PB2, PB1 and PA, 1.0 ng of NP, all in pCAGGS expression vector, as well as 20 ng pHH21-vNP-FF-Luc (firefly luciferase), 50 ng of pRL-SV40-R*luc* (Promega) and varying input (50 ng for 5-site screen and 15 ng for 4-site screen) of MxA plasmids were co-transfected into HEK 293T cells using the reagent TransIT-LT1 (Mirus Bio) in a 96-well plate format. After 24 h, firefly luciferase activity was measured using the Dual-Glo system (Promega) and normalized to the constitutively co-expressed *Renilla* luciferase (transfection efficiency control). Results are presented relative to the luciferase readings in the absence of MxA (‘empty vector’) as the average of fold restriction in three independent experiments. Expression of MxA proteins was detected by Western blot analysis using mouse ANTI-FLAG^®^ monoclonal antibody F4042 (Sigma) and rabbit anti-beta-actin polyclonal antibody ab8227 (Abcam) or anti-beta-tubulin ab6046 (Abcam). Corresponding data from minireplicon assays and Western blots are derived from samples generated from a single master mix. All experiments were done in triplicate.

For the IAV (H5N1) restriction analyses, we performed similar experiments but with the H5N1 minireplicon system of A/Vietnam/1203/04 as described (13) in a 12-well format, including a reporter construct encoding firefly luciferase under the control of the viral promoter, and 300 ng of the MxA expression plasmid. Just like with the THOV minireplicon experiments, restriction was assayed relative to empty vector control by measuring firefly luciferase expression relative to the *Renilla* luciferase transfection control.

### THOV infection assays

Huh7 cells were transfected with MxA expression plasmids (0.2 ng per 6-well) and infected 24 h later with THOV virus at an MOI of 5. After fixation of the cells 24 hours after infection, MxA and THOV NP were stained, and cells were analyzed by FACS. MxA-positive cells were selected after doublet exclusion, and the percentage of infected (THOV NP-positive) cells was determined. We first assessed what fraction of cells expressing the catalytically inactive MxA mutant T103A were infected by THOV (based on THOV NP expression) and set this fraction to 100%. We subsequently measured fraction of cells expressing different MxA variants that were infected by THOV to determine the amount of restriction provided by wtMxA, MxA super-restrictors, or MxA non-restrictors (PFFSS or ΔL4).

### Co-Immunoprecipitation

293T cells seeded in 6-well plates were transfected with 1 μg of expression plasmids coding for flag-tagged wtMxA and MxA variants using the JetPEI transfection reagent (Polyplus). 24 h post-transfection, cells were infected with THOV (MOI 10). 24 h post-infection, co-immunoprecipitation analysis was performed. The cells were lysed in 50 mM Tris (pH 8.0), 150 mM NaCl, 1 mM EDTA, 0.5% Nonidet P-40 and incubated with anti-FLAG-M2 affinity gel (Sigma-Aldrich) for 2 h at 4 °C. After extensive washing, the precipitates as well as the wholecell lysates were subjected to standard Western blot analysis using antibodies against the MxA protein and THOV NP as described previously (12).

### Logo plots

Difference logo plots comparing super-restrictors, restrictors or non-restrictor classes to the full library. For each class of clones, we constructed an amino acid position frequency matrix, and generated difference logo plots using the DiffLogo R package [37].

### Statistical Analysis

Data analyses were done using GraphPad Prism 7.0 software. All data are shown as mean ± SEM. Statistical analysis was performed using the non-parametric test Mann-Whitney U test (p values less than 0.05 were considered statistically significant). We used a Pearson’s correlation to measure the linear relationship between THOV restriction and IAV restriction for the various MxA super-restrictors and wtMxA.

## Acknowledgements

We thank HHMI EXROP interns María Gutierrez (UT El Paso) and Isabel Delwel (Univ. North Texas) for assistance with THOV and IAV minireplicon assays, and P. Mitchell and members of the Malik and Emerman labs for advice and discussions. We also thank J. Bloom, N. Elde, P. Mitchell, S. Soh, and J. Tenthorey for their comments on the manuscript.

## SUPPLEMENTARY FIGURE LEGENDS

**S1_Fig. Schematic of combinatorial mutagenesis screen.** The mutant library was generated using oligonucleotide-directed mutagenesis of all five rapidly-evolving sites in wtMxA L4 (Materials and Methods). The resulting gene library was cloned into the expression vector pQXCIP and transformed in bacterial cells. Six hundred individual colonies were randomly selected, plasmids were extracted and sequenced. The antiviral activity of each variant was evaluated on the THOV minireplicon assay (Materials and Methods) in triplicate. In a second iteration, we screened combinatorial mutants of L4 except for position 561 that was fixed as a Phenylalanine (F).

**S2_Fig. Calibrating the dynamic range of the minireplicon assay to measure MxA restriction.** We wished to maximize the sensitivity and dynamic range of the minireplicon assay to measure anti-THOV restriction activity of different MxA variants. For this, we transfected increasing amounts of wtMxA and the catalytically inactive mutant MxA-T103A expression plasmids with the THOV minireplicon into HEK-293T cells. The amounts used for the less-sensitive 5-site screen (50 ng) (Fig 1) and more sensitive 4-site screen (15 ng) (Fig 2) are indicated on the graph. Data is presented as a mean average of three experiments (error bars ±SEM).

**S3_Fig. Setting thresholds for wtMxA restriction levels for the 5-site MxA variant screen.** We tested 91 distinct clones of wtMxA and 13 MxA-T103A mutants in triplicates in the minireplicon assay. The range of restriction potential represents the wtMxA-equivalent range of restriction (dashed lines, Fig 1B).

**S4_Fig. Identifying amino acid residues enriched in the 4-site super-restrictor pool.** We used difference logo plots to compare super-restrictors to the full library. The stack height for each motif position is calculated using the Jensen-Shannon divergence (JS) (22). Amino acids shown above the y=0 line on the left are enriched in each class of clones compared to their background frequency in the whole library, and those above the y=0 line on the right are depleted. The total height of each stack of letters represents how different the two classes of clones are from one another, and the height of each amino acid letter reflects how much it contributes to the overall difference in frequencies. For example, Q540 is enriched in the super-restrictor pool in our analysis of 141 4-site MxA variants.

**S1_Table. THOV minireplicon measurements of restriction activity for 523 5-site MxA variants.** L4 sequence and ant-THOV restriction activity of individual MxA variants is reported.

**S2_Table. THOV minireplicon measurements of restriction activity for 141 4-site MxA variants.** L4 sequence and ant-THOV restriction activity of individual MxA variants is reported.

**S3_Table. Distribution of mutations.** Distribution of mutations relative to wtMxA in 4-site combinatorial mutagenesis screen of MxA L4.

**S4_Table. Numerical values.** Numerical values for all figures are reported.

**S1_Data. All original Western blots used in our analyses.**

## Notes

#### Summary of Updates

New experiments (Figure 3, Figure 5 C, D) revised Abstract, Results and Discussion; note author names corrected (middle initials) and new Author added

