## Supplementary figures and images for "Combinatorial mutagenesis of rapidly-evolving residues yields super-restrictor antiviral proteins"

### Supplementary Figure 1

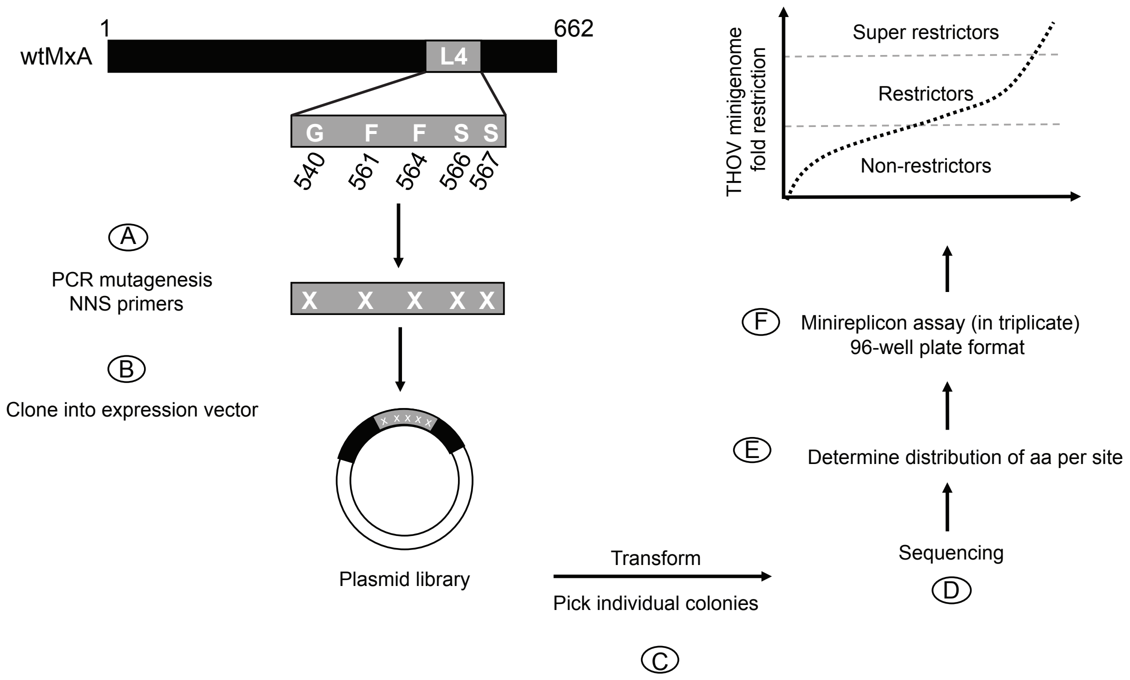

### Supplementary Figure 2

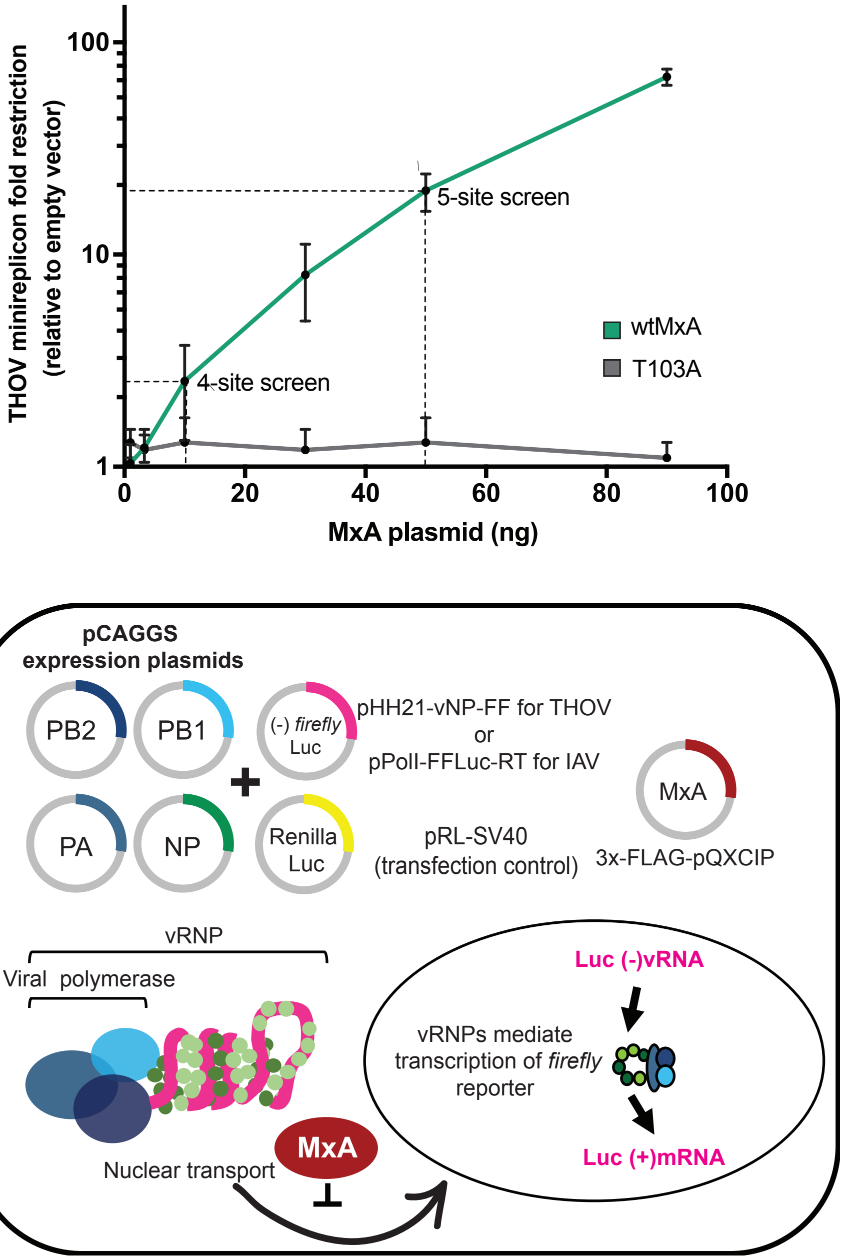

### Supplementary Figure 3

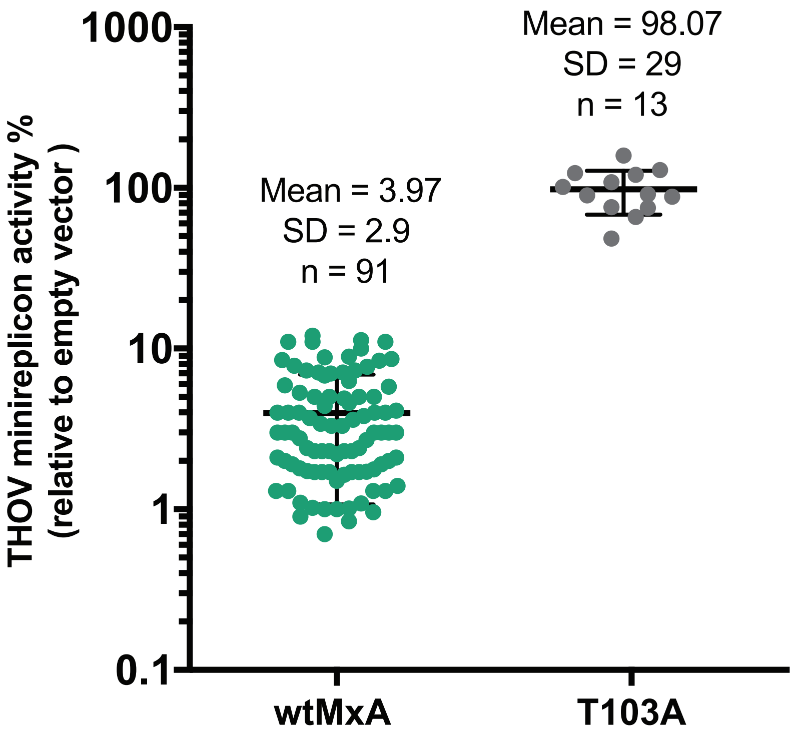

### Supplementary Figure 4

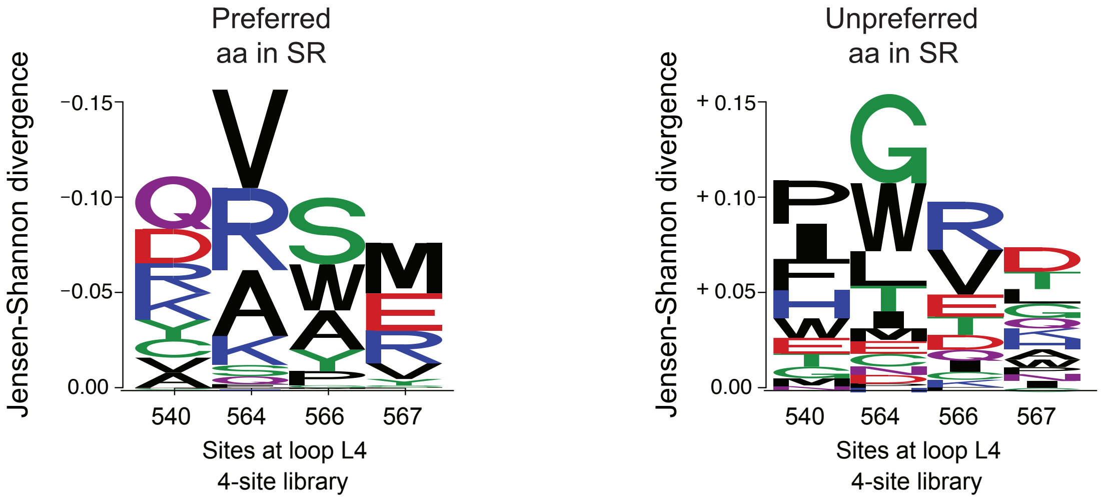
